## Supplemental Data for "Maf-family bZIP transcription factor NRL interacts with RNA-binding proteins and R-loops in retinal photoreceptors"

#### Supplemental Material

**Table S1. PSM scores of NRL-interacting retinal proteins identified by Liquid Chromatography with Tandem Mass Spectrometry (LC MS-MS) after purification with NRL-GST.** PSM values are shown for each protein. The listed proteins were enriched in three independent experiments.

| Accession | Gene | Protein name | GST 1 | NRL 1 | GST 2 | NRL 2 | GST 3 | NRL 3 |
|---|---|---|---|---|---|---|---|---|
| XP_005208934.1 | HNRNPM | Heterogeneous nuclear ribonucleoprotein M isoform X11 [Bos taurus] | | 68 | 1 | 163 | 4 | 109 |
| AAI33330.1 | HNRNPU | Heterogeneous nuclear ribonucleoprotein U (scaffold attachment factor A) [Bos taurus] | 1 | 56 | | 98 | 2 | 62 |
| DAA13799.1 | HNRNPUI1 | TPA: heterogeneous nuclear ribonucleoprotein U-like [Bos taurus] | | 9 | | 13 | | 9 |
| 122145945 | HNRNPA2/B1 | Heterogeneous nuclear ribonucleoproteins A2/B1 | | 4 | | 26 | | 17 |
| 82592672 | CFDP2 | Craniofacial development protein 2 | | 6 | | 18 | | 11 |
| 122143188 | PFKM | 6-phosphofructokinase, muscle type | | 2 | | 17 | | 12 |
| 2500541 | DHX9 | ATP-dependent RNA helicase A | | 5 | | 16 | | 5 |
| 172046785 | HNRNPA1 | Heterogeneous nuclear ribonucleoprotein A1 | | 1 | | 16 | | 12 |
| 122142416 | AP2A2 | AP-2 complex subunit alpha-2 | | 1 | | 9 | | 6 |
| 1169457 | EAAT1 | Excitatory amino acid transporter 1 | | 1 | | 9 | | 8 |
| 399012 | SLC25A6 | ADP/ATP translocase 3 | | 1 | | 6 | | 2 |
| 109892458 | HNRNPH2 | Heterogeneous nuclear | | 1 | | 5 | | 3 |

| | | | | | | | | |
|---|---|---|---|---|---|---|---|---|
| | | ribonucleoprotein H2 | | | | | | |
| 122132446 | STRBP | Spermatid perinuclear RNA-binding protein | | 3 | | 5 | | 3 |
| 75057558 | EPB41L5 | Band 4.1-like protein 5 | | 1 | | 4 | | 3 |
| 547891 | MAP4 | Microtubule-associated protein 4 | | 1 | | 4 | | 4 |
| 108860929 | RPL23A | 60S ribosomal protein L23a | | 1 | | 3 | | 2 |
| 2495339 | HSPA1B | Heat shock 70 kDa protein 1B | | 1 | | 3 | | 3 |
| 54040030 | YBX1 | Nuclease-sensitive element-binding protein 1 | | 1 | | 3 | | 2 |
| 122132456 | PRPF19 | Pre-mRNA-processing factor 19 | | 1 | | 3 | | 3 |
| 129204 | RHO | Rhodopsin | | 1 | | 3 | | 1 |
| 166219437 | RPS17 | 40S ribosomal protein S17 | | 1 | | 2 | | 1 |
| 387935413 | ZNF326 | DBIRD complex subunit ZNF326 | | 1 | | 2 | | 1 |
| 110278997 | HIST1H2BK | Histone H2B type 1-K | | 1 | | 2 | | 3 |
| 126664 | SLC25A11 | Mitochondrial 2-oxoglutarate/malate carrier protein | | 2 | | 2 | | 2 |
| 121750 | SLC2A1 | Solute carrier family 2, facilitated glucose transporter member 1 | | 1 | | 2 | | 1 |
| 120752 | GABRA1 | Gamma-aminobutyric acid receptor subunit alpha-1 | | 2 | | 1 | | 1 |
| 122143023 | H2AFJ | Histone H2A.J | | 1 | | 1 | | 1 |
| 148887198 | HSPA8 | Heat shock cognate 71 kDa protein | | 6 | 1 | 13 | | 8 |
| 1709348 | NRL | Neural retina-specific leucine zipper protein | 1 | 15 | 4 | 36 | 4 | 32 |

Figure S1

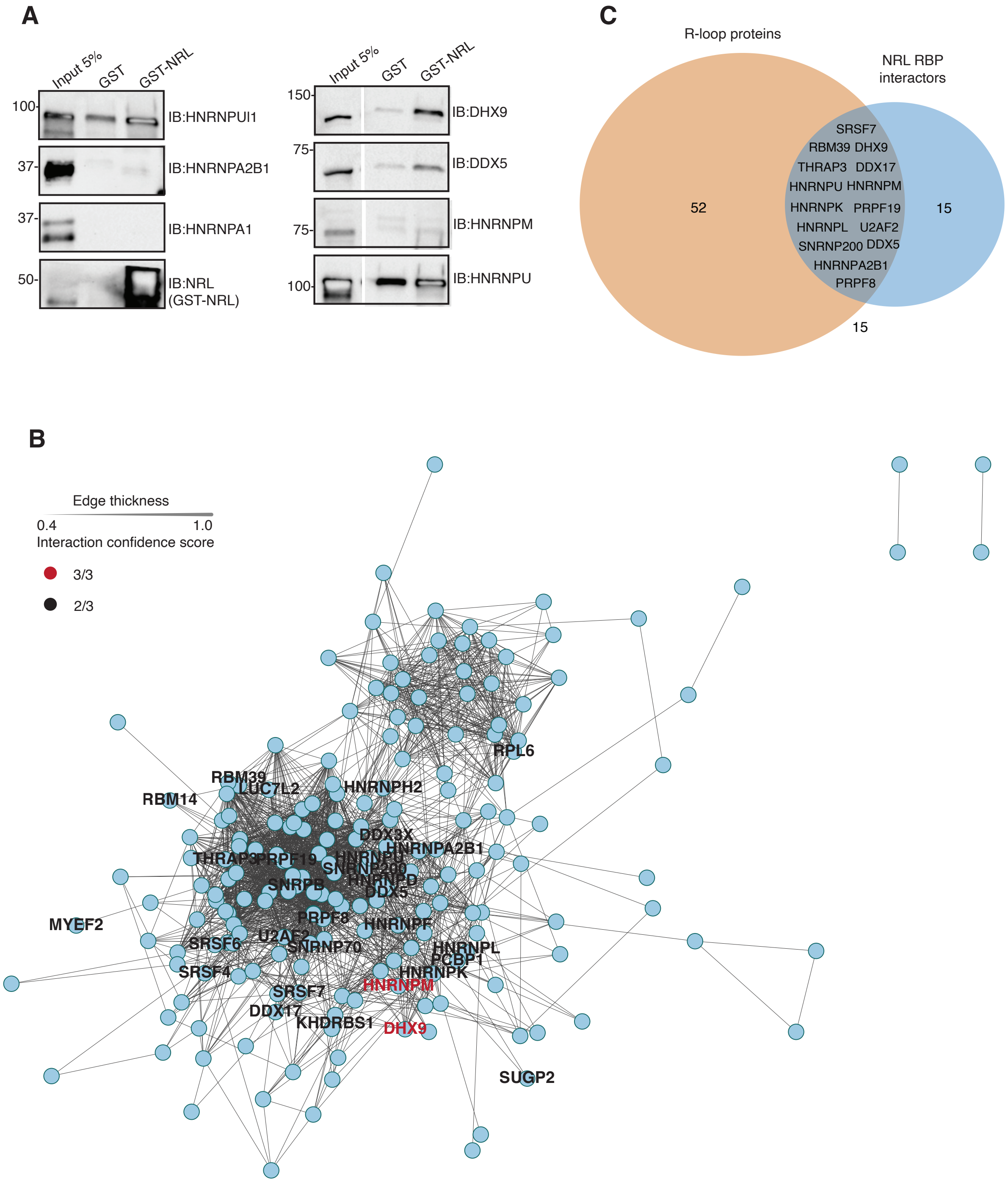


**Figure S1. NRL-RBP interactors are enriched in R-loop proteins.** A) Western blot of GST-NRL affinity-purified RBPs from bovine retina. Benzonase was added to retinal lysates during GST-NRL incubations. B) PPI network showing RBP experimental interactions from String. The edge thickness represents the confidence score with a cutoff of 0.4. Name of proteins identified in 2/4 assays are shown. Proteins identified in 3 out of 4 assays are highlighted in red. C) Venn diagram showing intersection of R-loop proteomes common to four studies [41, 42, 43, 44] and NRL RBP candidate interactors identified in 2/4 assays.

Figure S2

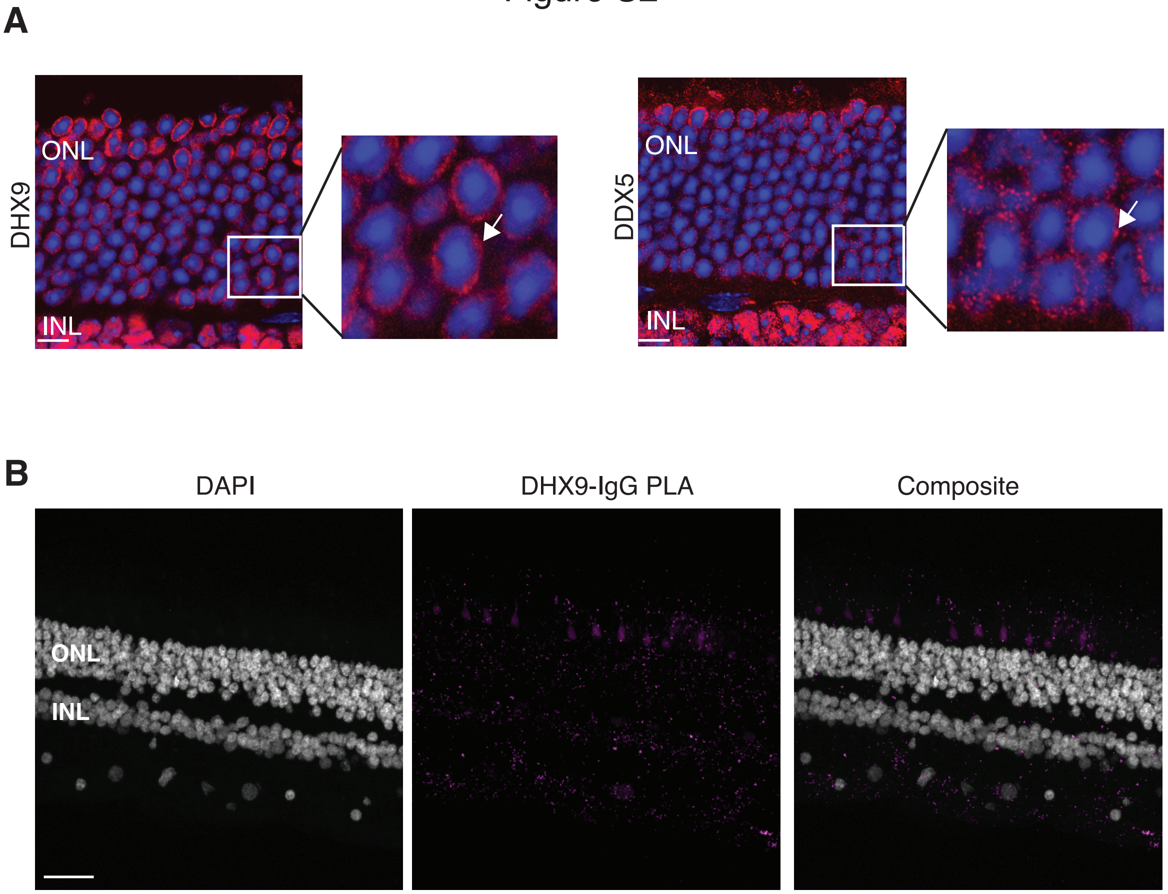


**Figure S2. Subcellular localization of DHX9 and DDX5 in adult mouse retina (P28).** A) DHX9 and DDX5 localization to the euchromatin region (nuclear periphery in murine rods) is shown in red. Zoom-in insets show proteins distributed in puncta (arrows). Nuclei is stained with DAPI. Scale bar is 10 μM. B) Proximity ligation assay (PLA) signal (magenta) using anti-DHX9 and goat IgG antibodies in the adult human retina. Scale bar = 20 μM. ONL = Outer nuclear layer; INL = Inner nuclear layer.

### Figure S3

**A**

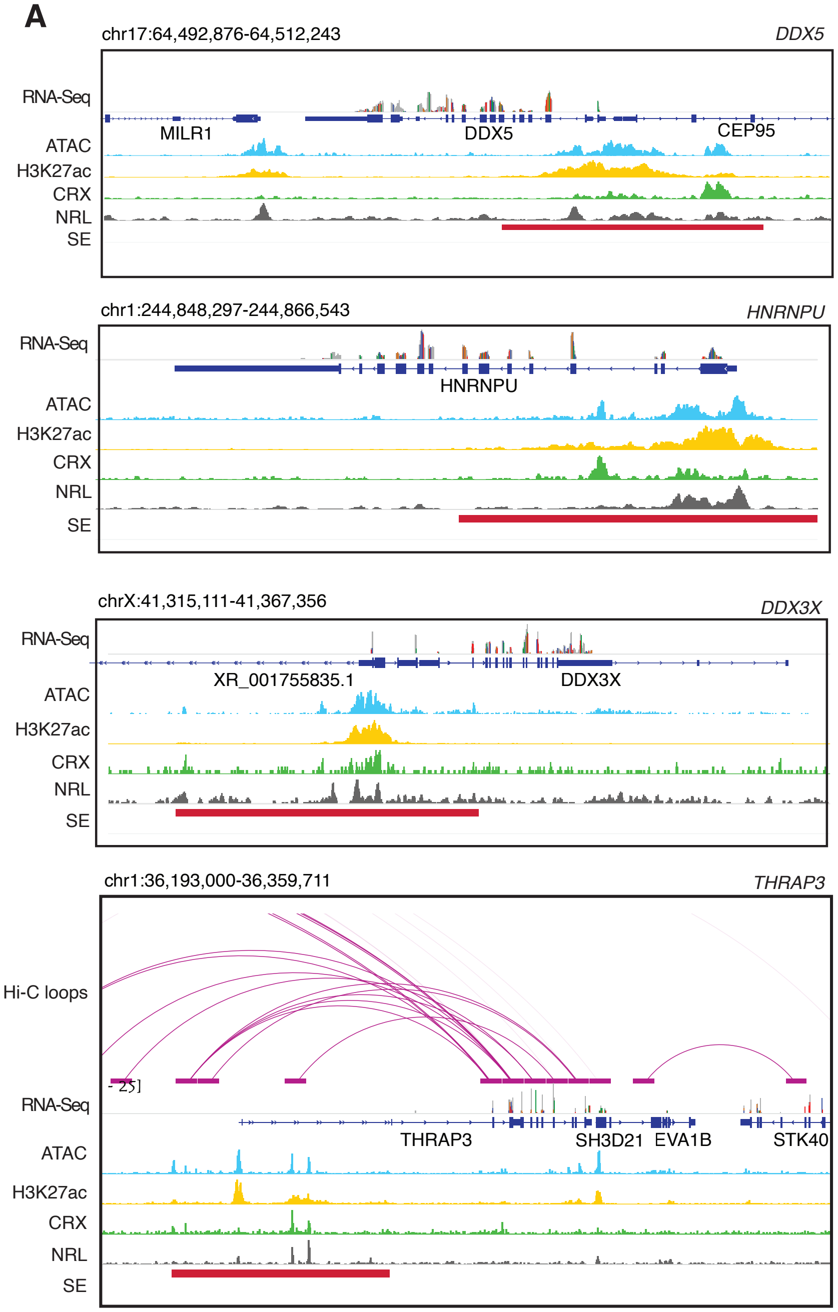
chr17:64,492,876-64,512,243
DDX5
RNA-Seq
MILR1
DDX5
CEP95
ATAC
H3K27ac
CRX
NRL
SE
chr1:244,848,297-244,866,543
HNRNPU
RNA-Seq
HNRNPU
ATAC
H3K27ac
CRX
NRL
SE
chrX:41,315,111-41,367,356
DDX3X
RNA-Seq
XR_001755835.1
DDX3X
ATAC
H3K27ac
CRX
NRL
SE
chr1:36,193,000-36,359,711
THRAP3
Hi-C loops
RNA-Seq
THRAP3
SH3D21
EVA1B
STK40
ATAC
H3K27ac
CRX
NRL
SE

**B**

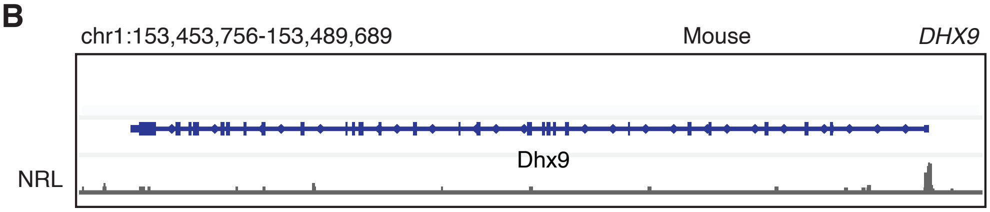
chr1:153,453,756-153,489,689
Mouse
DHX9
Dhx9
NRL

**Figure S3. NRL occupancy on super enhancers at genes encoding NRL-interacting RBPs.** A) Genomic view of the human *DDX5, HNRNPU, DDX3X, and THRAP* loci showing Hi-C loops, RNA-Seq, ATAC-Seq, H3K27ac ChIP-Seq, CRX-ChIP-Seq, NRL-ChIP-Seq and super-enhancer tracks (Obtained from *Marchal et al. 2022*). B) Genome browser view showing Cut&Run peaks for NRL at mouse *Dhx9* promoter.

Figure S4

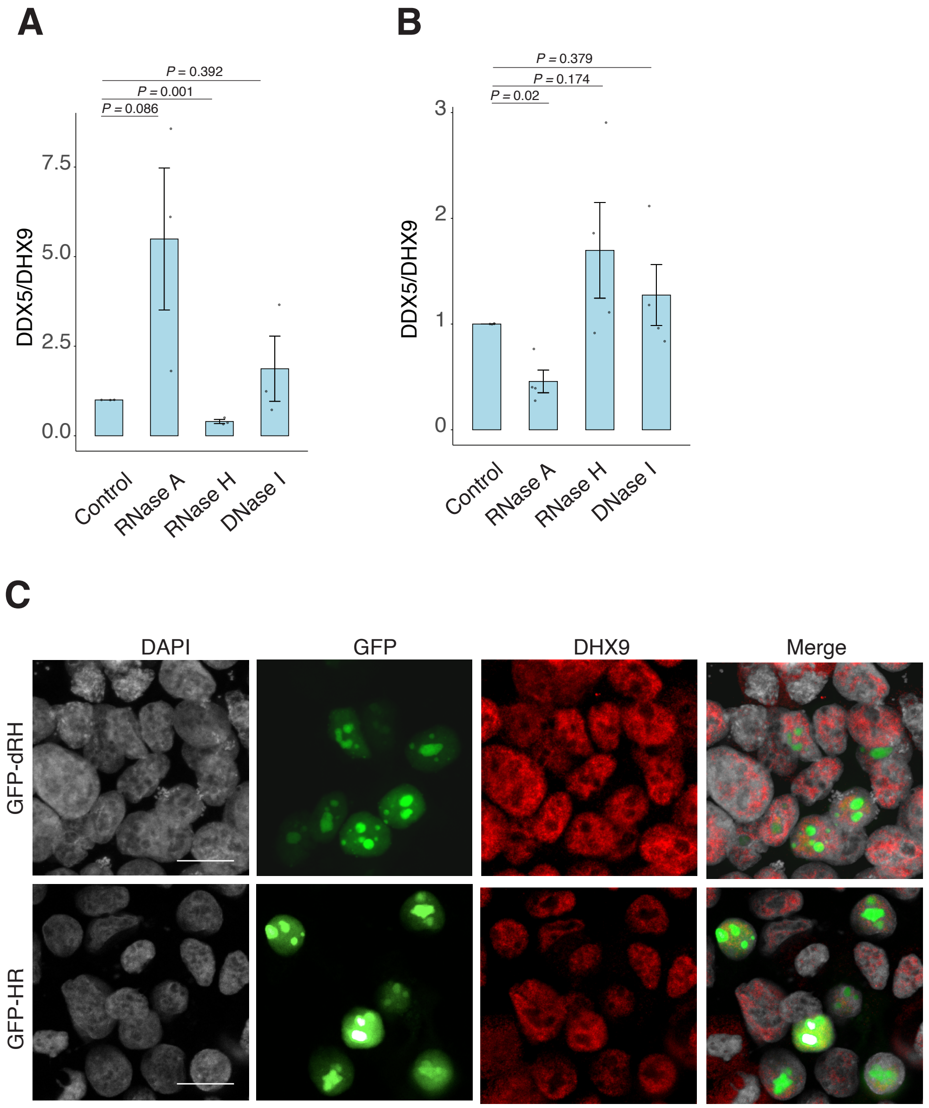


**Figure S4. DHX9 interacts with DDX5 in HEK293 and bovine retina.** A-B) Co-immunoprecipitation (co-IP) of DDX5 with DHX9 antibody in HEK293 cells overexpressing NRL (A) and in bovine retina (B). Lysates were treated for 30 min with different nucleases (as shown) before incubations with DHX9 antibody (n = 3 for HEK cells; n = 4 for bovine retina). Quantification of signal intensities were normalized to precipitated DHX9. Data are presented as the mean ± SEM. Unpaired two-tailed t test was performed to compare means of samples against controls. C) Confocal images of cells transfected with NRL and wild type (WT) human RNase H1 or D201N catalytic dead mutant EGFP fusions (GFP-HR, and GFP-dHR, respectively). Cells were stained with antibodies against DHX9 (red). Nuclei were stained with DAPI (grey). Scale bar is 20 μM.

Figure S5

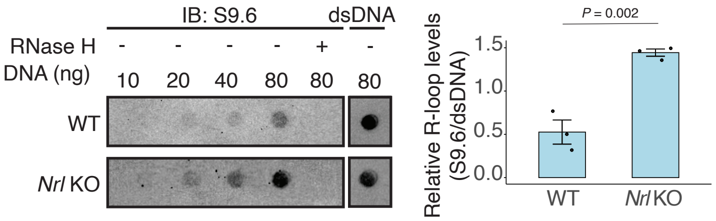


**Figure S5. R-loops are increased in NRL KO retina.** Dot blot of DNA:RNA hybrids from adult wild type and Nrl KO retina. Genomic DNA (gDNA) from retina was treated with RNaseIII with and without RNase H overnight. R-loops were detected using S9.6 antibody. Bar graph shows quantification of R-loop levels in wild type compared to *Nrl* KO retina (n = 3). Data are presented as the mean ± SEM. Unpaired two-tailed t test was performed to compare means of samples against controls.

### Figure S6

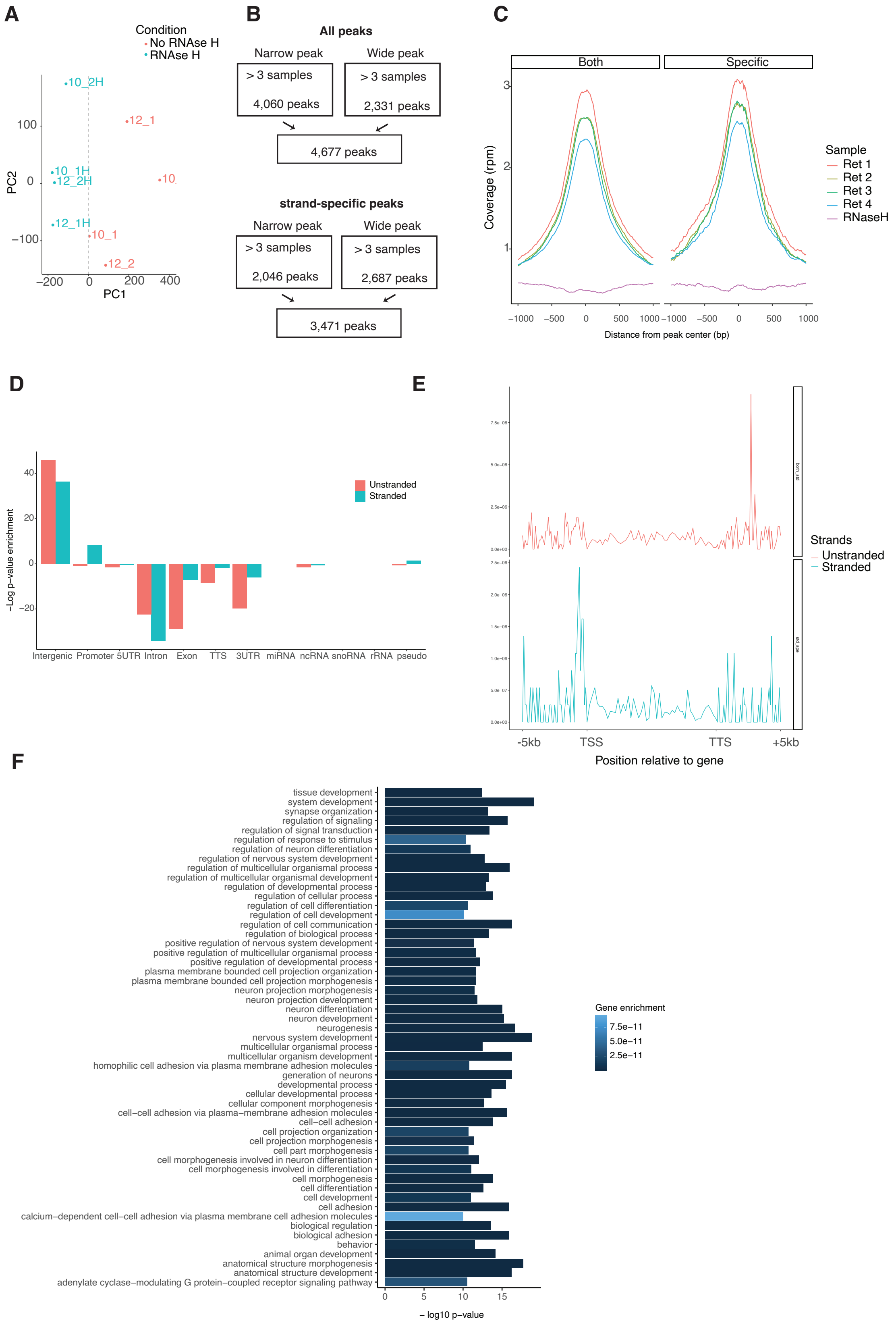

A
Condition
No RNAse H
RNAse H
PC2
PC1
B
All peaks
Narrow peak
Wide peak
> 3 samples
4,060 peaks
> 3 samples
2,331 peaks
4,677 peaks
strand-specific peaks
Narrow peak
Wide peak
> 3 samples
2,046 peaks
> 3 samples
2,687 peaks
3,471 peaks
C
Both
Specific
Coverage (rpm)
Distance from peak center (bp)
Sample
Ret 1
Ret 2
Ret 3
Ret 4
RNaseH
D
−Log p−value enrichment
Unstranded
Stranded
Intergenic Promoter 5UTR Intron Exon TTS 3UTR miRNA ncRNA snoRNA rRNA pseudo
E
Strands
Unstranded
Stranded
-5kb TSS TTS +5kb
Position relative to gene
F
tissue development
system development
synapse organization
regulation of signaling
regulation of signal transduction
regulation of response to stimulus
regulation of neuron differentiation
regulation of nervous system development
regulation of multicellular organismal process
regulation of multicellular organismal development
regulation of developmental process
regulation of cellular process
regulation of cell differentiation
regulation of cell development
regulation of cell communication
regulation of biological process
positive regulation of nervous system development
positive regulation of multicellular organismal process
positive regulation of developmental process
plasma membrane bounded cell projection organization
plasma membrane bounded cell projection morphogenesis
neuron projection morphogenesis
neuron projection development
neuron differentiation
neuron development
neurogenesis
nervous system development
multicellular organismal process
multicellular organism development
homophilic cell adhesion via plasma membrane adhesion molecules
generation of neurons
developmental process
cellular developmental process
cellular component morphogenesis
cell−cell adhesion via plasma−membrane adhesion molecules
cell−cell adhesion
cell projection organization
cell projection morphogenesis
cell part morphogenesis
cell morphogenesis involved in neuron differentiation
cell morphogenesis involved in differentiation
cell morphogenesis
cell differentiation
cell development
cell adhesion
calcium−dependent cell−cell adhesion via plasma membrane cell adhesion molecules
biological regulation
biological adhesion
behavior
animal organ development
anatomical structure morphogenesis
anatomical structure development
adenylate cyclase−modulating G protein−coupled receptor signaling pathway
− log10 p−value
Gene enrichment
7.5e−11
5.0e−11
2.5e−11

**Figure S6. Epigenomic signatures of R-loop formation in the retina.** A) Principal components analysis on scaled DRIP-seq coverage (500bp windows). PC1 (x axis) and PC2 (y axis) are plotted. Color represents samples treated (blue) or not (red) with RNase H. The grey dotted line shows the PC1 value separating treated vs non-treated samples. B) R-loop peaks from ssDRIP-Seq identified with narrow or broad peak parameters using RNase H treated sample (upper flow chart) or the opposite strand (lower flow chart) for enrichment. C) Metaplot showing coverage per ssDRIP sample on stranded and unstranded R-loop peaks. D) Metaplot of ssDRIP-seq signals for stranded or unstranded R-loops centered on gene bodies +/-5 kb. E) Enrichment of unstranded and stranded R-loops at different genomic regions. F) Biological process enrichment of genes associated with stranded R-loops.

### Figure S7

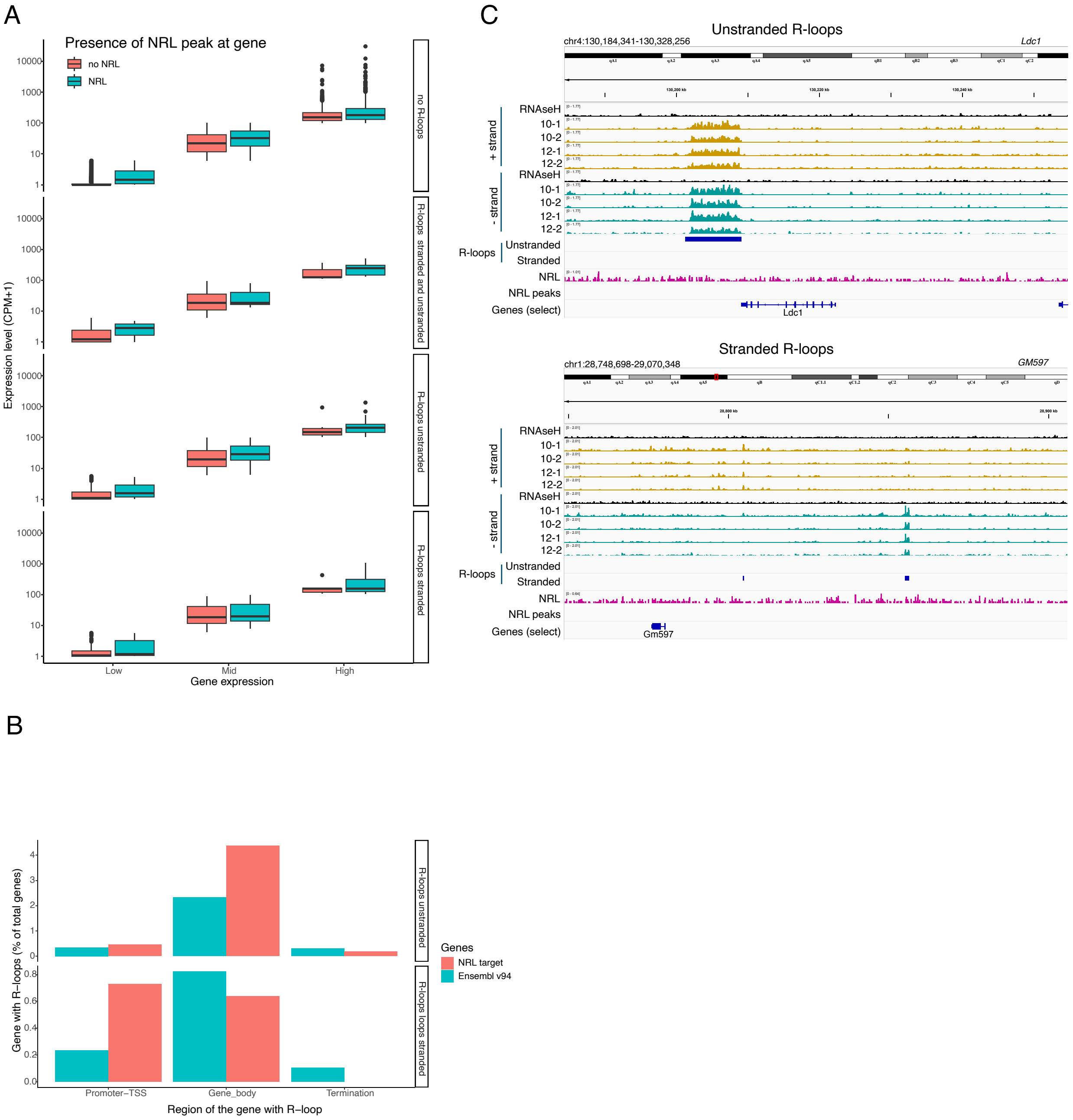


**Figure S7. Distribution of NRL peaks over genes.** A) Boxplot showing the expression level of low, mid and highly expressed genes with or without NRL binding. B) Bargraph showing R-loops over different gene regions at all genes (Ensemble v94) and genes regulated by NRL (Obtained from Liang et al., 2022). C) Genome view of *Ldc1* mouse gene and an intergenic region displaying ssDRIP-Seq signal in four retinas. Signals are shown for the positive (orange) and negative (blue) strands separately. RNase H treated samples are pooled and shown for each strand (black). Peak calls for NRL and unstranded and stranded R-loops are shown in blue.

Figure S8

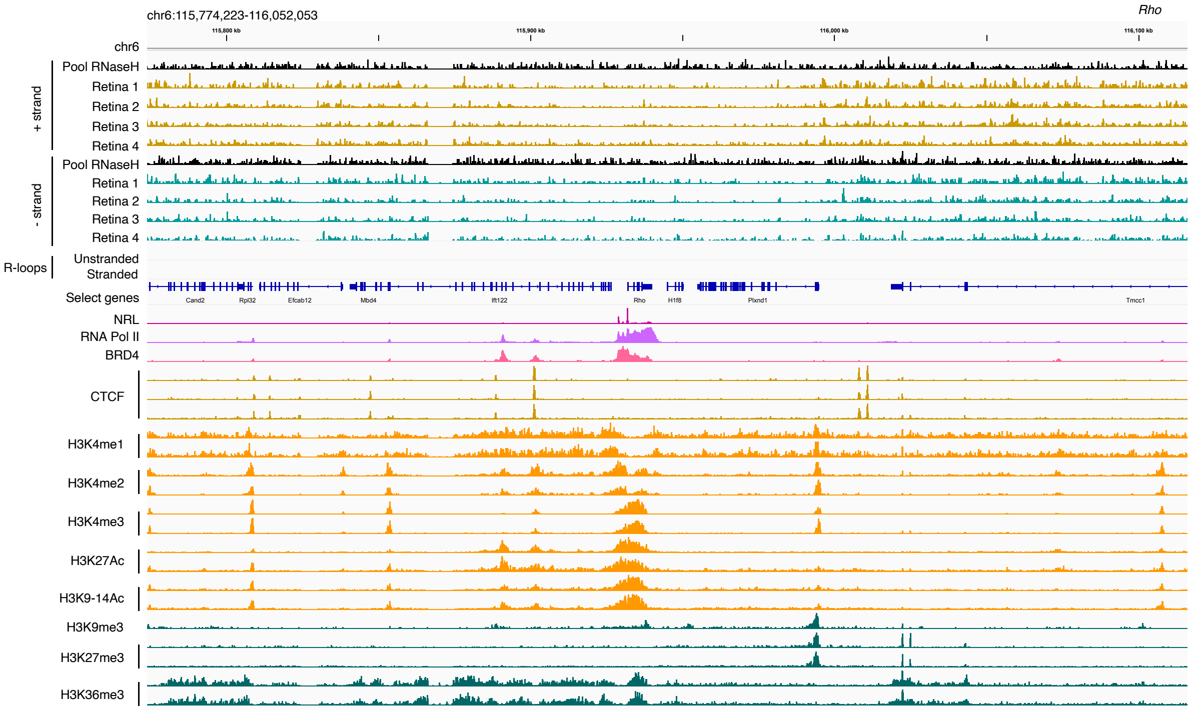


**Figure S8. Absence of R-loops over the *Rho* gene.** Genome view of *Rho* mouse gene and neighboring loci displaying ssDRIP-Seq signal in four retinas. Signals are shown for the positive (orange) and negative (blue) strands separately. RNase H-treated samples are pooled and shown for each strand (black). Peak calls for NRL and unstranded and stranded R-loops are not present. Signals for NRL, RNA pol II, BRD4, CTCF and various histone modifications are shown.
